# Transcriptomic analysis of diplomonad parasites reveals a trans-spliced intron in a helicase gene in *Giardia*

**DOI:** 10.1101/052134

**Authors:** Scott William Roy

## Abstract

Gene expression is the central preoccupation of molecular biology, thus newly discovered facets of gene expression are of great interest. Recently, ourselves and others reported that in the diplomonad protist *Giardia lamblia*, the coding regions of several mRNAs are produced by ligation of independent RNA species expressed from distinct genomic loci. Such trans-splicing of introns was found to affect nearly as many genes in this organism as does classical cis-splicing of introns. These findings raised questions about the incidence of intron trans-splicing both across the *G. lamblia* transcriptome and across diplomonad diversity, however a dearth of transcriptomic data at the time prohibited systematic study of these questions. Here, I leverage newly available transcriptomic data from *G. lamblia* and the related diplomonad *Spironucleus salmonicida* to search for trans-spliced introns. My computational pipeline recovers all four previously reported trans-spliced introns in *G. lamblia*, suggesting good sensitivity. Scrutiny of thousands of potential cases revealed only a single additional trans-spliced intron in *G. lamblia*, in the p68 helicase gene, and no cases in *S. salmonicida*. The p68 intron differs from the previously reported trans-spliced introns in its high degree of streamlining: the core features of *G. lamblia* trans-spliced introns closely packed together, revealing striking efficiency in the implementation of a seemingly inherently inefficient molecular mechanism. These results serve to circumscribe the role of trans-splicing both in terms of genes effected and taxonomically. Future work should focus on the molecular mechanisms, evolutionary origins and phenotypic implications of this intriguing phenomenon.

---

Splicing of nuclear RNA transcripts by the spliceosomal machinery is a ubiquitous feature of the expression of nuclear genes in eukaryotes (Roy and Irimia 2014; Nixon et al. 2002; Vanacova et al. 2005; although see Lane et al. 2007; Akiyoshi et al. 2009). Splicing within protein-coding sequences nearly always joins two protein-coding regions of a single RNA transcribed from a single locus: intron *cis-*splicing (Chow et al. 1977). Alternatively, protein-coding regions from multiple RNAs transcribed from different loci can be joined: intron *trans*-splicing (Li et al. 1999; Takahara et al. 2000; Dorn et al. 2001; Robertson et al. 2007; Fang et al. 2012). (This process should be distinguished from spliced leader trans-splicing, in which a short non-coding RNA molecule is added to various mRNAs outside of the coding region, essentially donating 5 UTR sequence (Lasda and Blumenthal 2011). *Trans*-splicing of introns is generally very rare: for instance, among the hundreds of thousands of known splicing events in humans, there are fewer than 10 confirmed cases of genic trans-splicing (Wu et al. 2014). Recently, the first case in which a substantial of introns in an organism are *trans*-spliced was reported. In the genome of the diplomonad intestinal parasite *G. lamblia*, systematic studies have revealed only six *cis*-spliced introns to date (Nixon et al. 2002; Russell et al. 2005; Morrison et al. 2007; Roy et al. 2012; Franzen et al. 2013); strikingly, small-scale studies revealed four cases of genic trans-splicing, including two in a single gene (Nageshan et al. 2011; Kamikawa et al. 2011; Roy et al. 2012; Hudson et al. 2015). These cases showed intriguing sequence features - most notably extended basepairing potential between the pairs of trans-spliced loci.

These studies raised two important questions. First, given the fact that these cases were found largely serendipitously, with a single gene containing two separate trans-spliced introns, is genic trans-splicing in *G. lamblia* much more widespread? Second, what is the evolutionary history of trans-splicing in *G. lamblia* and other diplomonads? However, the lack of availability of large amounts of mRNA sequence data at that time prohibited systematic study of these questions. Recently, Franzen et al. (2013) reported a transcriptome analysis of three different strains of *G. lamblia* and Xu et al. (2014) reported the genome and transcriptome of the distantly-related diplomonad parasite *Spironucleus salmonicida*. Here, I report the first transcriptome-wide studies of intron *trans*-splicing in *G. lamblia* isolates and *S. salmonicida*.

## Transcriptomic analysis of trans-splicing in diplomonad parasites

I downloaded 11 Illumina RNA-seq datasets from previous transcriptomic analyses, 10 for *G. lamblia* parasites from Franzen et al. (2013) and one of *S. salmonicida* from Xu et al. (2014). For each species, I used bowtie and blat to identify Illumina reads that contained sequence from multiple genomic loci and which are suggestive of *trans*-splicing (see Methods). This procedure identified some 495,066 potential boundaries in *G. lamblia* and 231,769 in *S. salmonicida*. For both species, the vast majority of these cases were either supported by only a single read (400,460 and 212,801 respectively), had extended similarities at the 5′ and 3′ boundaries suggesting reverse transcriptase artifacts produced during library formation (‘RTfacts’; Roy and Irimia 2008) (388,835 and 159,836 cases), and/or did not represent a clear splice junction (with >5 nucleotides in the middle of the read that did not map to either locus (35,740 and 8307cases). Filtering of these dubious cases left 2272 potential boundaries in *G. lamblia* and 5454 in *S. salmonicida*.

All 2272 *G. lamblia* cases and the most promising 500 *S. salmonicida* cases (see Methods) were analyzed by eye for presence of sequences corresponding to extended 5′ or 3′ splicing signals particular to the species. In *G. lamblia*, this analysis yielded five clear cases in *G. lamblia* and no “borderline” cases. That is, each of the five cases had extended an extended 5′ splicing signal (consensus GTATGTT) an extended 3′ splicing signal (CT[AG]ACACACAG), complementarity between the pairs of apparently trans-spliced loci, and presence of the *G. lamblia* 3 cleavage motif (consensus sequence TCCTTTACTCAA); no other cases showed any of these features. To confirm this manual analysis, all potential boundaries were also analyzed for adherence of splicing motifs to those of all known 10 *cis*- and *trans*-spliced introns (10 total in *G. lamblia*, 4 total in *S. salmonicida*), using a position weight matrix (PWM) approach (Figure 1a). These automated analyses confirmed the findings of only a single new case that exhibited canonical splicing boundaries. A similar combination of manual and automated PWM analysis in *S. salmonicida* did not yield any strong trans-splicing candidates (Figure 1b).

**Figure 1.**
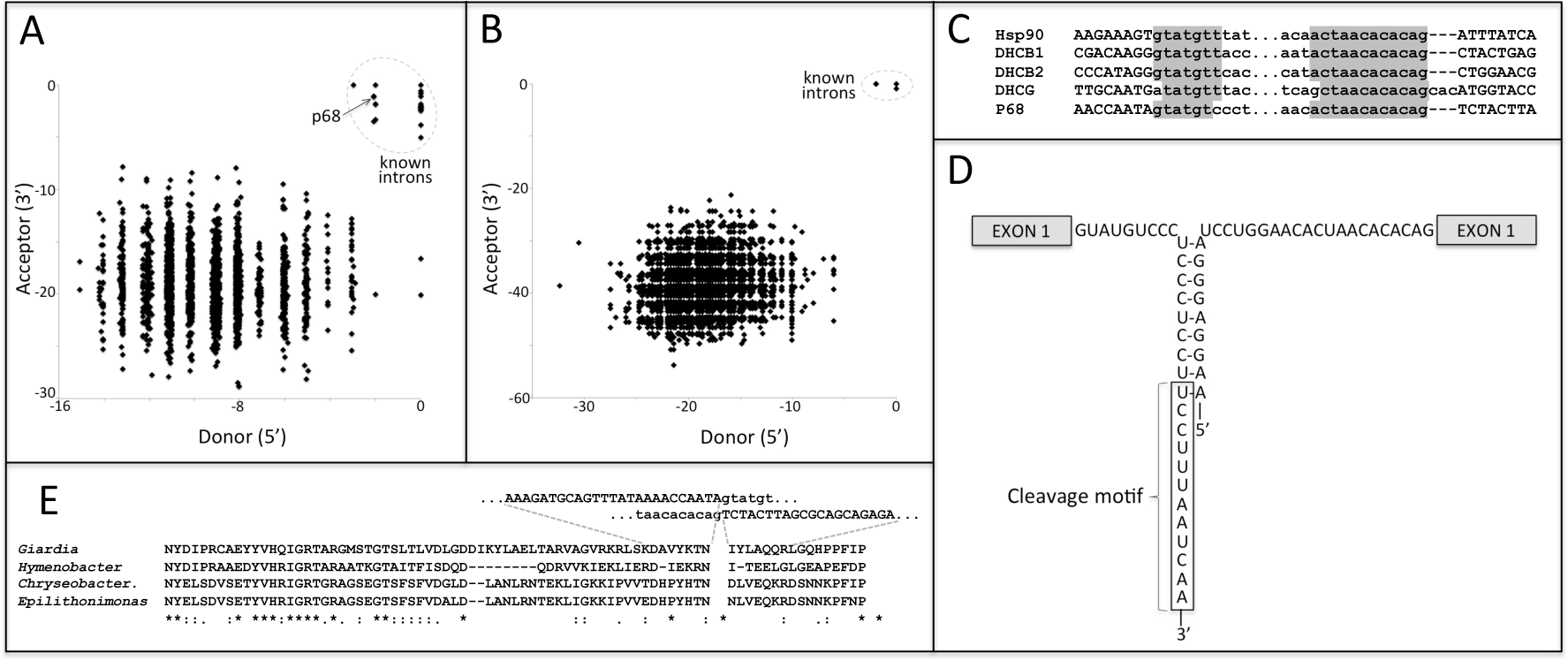
Transcriptome-wide search identifies a trans-spliced intron in p68 helicase. A. Normalized PWM scores for similarity to known donor and acceptor intron splice boundaries for 2272 potential *G. lamblia* trans-splicing events supported by at least two reads and for known *cis-* and trans-spliced introns. Only one potential trans-splicing events, in the p68 helicase, groups with known introns. Boundaries are scored relative to the maximum possible score (equating to zero). B. Normalized PWM scores for similarity to known donor and acceptor intron splice boundaries for 5454 potential *S. salmonicida* trans-splicing events supported by at least two reads and for known c/s-spliced introns. No potential trans-splicing events group with known introns. C. Comparison of splice boundaries for newly-discovered p68 trans-spliced intron with other trans-spliced introns, for *G. lamblia* isolate GS. D. Trans-spliced intron sequences for newly-discovered p68 intron exhibits basepairing potential between intronic regions of 5’ and 3’ pre-mRNA transcripts and conserved cleavage motif reported by Hudson et al. (2012). E. Protein sequence alignment between the protein encoded by trans-spliced *G. lamblia* p68 gene and closest BLAST hits in Genbank *[Chryseobacterium caeni*, Accession WP_027384510.1, *Hymenobacter sp. AT01-02*, Accession WP_052694982.1, and *Epilithonimonas tenax*, Accession WP_028122041.1).

## A trans-spliced intron in a p68 helicasegene

I next focused on the five identified trans-splicing candidates in *G. lamblia*. Mapping of RNA-seq data from Franzen et al. (2013) and from a second study that became available during the course of this project (Ansell et al. 2015) revealed direct RNA support for all five of these trans-splicing events across various *G. lamblia* isolates and conditions, with between 388 and 55,177 total reads supporting the five cases (Table 1). Four of these cases corresponded to all four previously reported trans-spliced introns. The fifth case represents a previously-unreported case of trans-splicing, falling in a putative p68 RNA-dependent helicase gene.

**Table 1.**
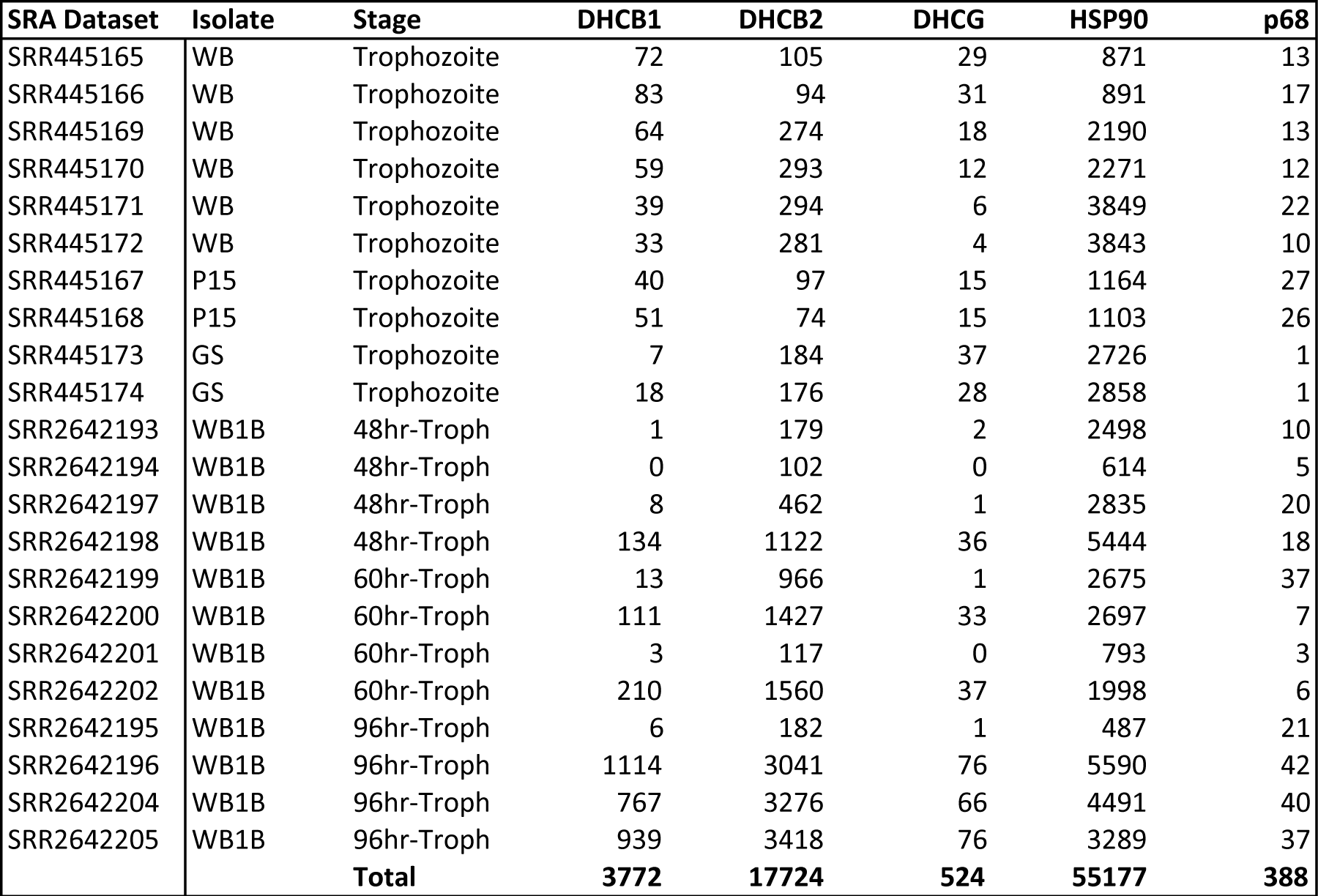
Number of reads supporting trans-splicing of five trans-spliced *G. lamblia* introns from mixed stage or synchronized stage trophozoites from 22 Illumina RNA-seq datasets. 48hr/60hr/96hr-Troph indicate hours after beginning of the trophozoite stage (for details see Ansell et al. (2015)). DHCB1/2: first/second intron of dynein heavy chain beta. DHCG: dynein heavy chain gamma.

This new trans-spliced intron exhibits the characteristic traits of the four previously-reported trans-spliced introns: (i) extended 5′ and 3′ splice sites (GTATGT and ACTAACACAG, respectively; Figure 1c); (ii) extended basepairing between the intronic regions of the two pre-mRNA transcripts (Figure 1d); and (iii) the recently-discovered *G. lamblia* cleavage motif (with consensus TCCTTTACTCAA; Figure 1d; Hudson et al. 2012). A BLASTX search of the mature trans-spliced transcript against Genbank revealed homology to p68 helicase (Figure 1e). Interestingly, the best several Genbank hits were bacterial, suggesting the possibility of lateral transfer of this gene to *G. lamblia*, and thus suggesting that one of the few introns in *G. lamblia* is a relatively new acquisition. This newly discovered intron represents an apex of structural economy among trans-spliced introns, with a short stretch of perfect Watson-Crick basepairing directly followed by (indeed, overlapping) the cleavage motif (Figure 1d). For comparison, the cleavage motif in the p68 intron lies only 17 nucleotides downstream of the 5′ splice site, compared to 34-93 nucleotides in the four previously-described *trans*-spliced introns.

## The extent of intron trans-splicing in time and space

While the available test set is regrettably small, the finding that our transcriptomic pipeline was able to identify all four previously reported cases of *G. lamblia* trans-splicing suggests that the pipeline does not have a terribly high false negative rate. As such, that the pipeline identified only a single additional case of trans-splicing suggests that the breadth of trans-splicing within the *G. lamblia* transcriptome may be limited. Similarly, that the pipeline did not identify promising trans-splicing candidates in *S. salmonicida* further suggests that the phylogenetic breadth of trans-splicing within diplomonads may similarly be limited (consistent with the findings of Xu et al. 20014). Future work should focus on better understanding the diversity and origins of trans-splicing within relatives of *G. lamblia*.

## Concluding remarks

These results enrich the set of known trans-spliced introns in *G. lamblia* while at the same time circumscribing the likely transcriptome-wide importance of trans-splicing in this organism. The structural economy of the reported p68 helicase intron reveals a remarkable efficiency in implementing the seemingly inherently inefficient molecular mechanism of trans-splicing. These cases together represent a further embellishment on the core mechanisms of gene expression. As with previously described embellishments - intron splicing, alternative splicing and promoter usage, spliced leader trans-splicing, ribosomal readthrough and frameshifting, etc. - attention now turns to understanding the mechanisms, evolutionary origins and potential phenotypic implications of these intriguing trans-spliced introns.

## Methods

Full genome sequences and Illumina RNA-seq data were downloaded for three strains of *G. lamblia* (GEO accession GSE36490, from Franzen et al. 2013) and for *S. salmonicida* (SRA accession SRR948595, from Xu et al. 2014). Bowtie (Langmead et al. 2009) was used with default parameters to exclude read pairs that mapped in expected orientation to the genome (with a maximum insert size 1000 nucleotides) and as well as individual reads that mapped to the genome. I then mapped the non-mapping reads to the genome using blat (Kent 2002) and identified reads for which (i) parts of the read mapped in exactly two places; (ii) both the 5′ and 3′ termini of the read mapped (that is, the mapping started within 5 nucleotides of the end of the read); and (iii) the junction between the two mappings was relatively precise-single unambiguous junction of with five or fewer nucleotides of overlap (i.e., in cases of similarity between the genomic sequences at the boundaries of the junction) or of gap (i.e., nucleotides near the junction that are not represented in either genomic locus). Junctions supported by at least two reads that suggested trans-splicing (either >5kb apart on the same contig or on different contigs) were then collected.

Each potential case of trans-splicing was assigned a 5′ and 3′ score based on adherence to splice boundaries of known introns. Scores were calculated using a standard PWM approach as follows: (i) 5′ and 3′ splice sites were compiled for all known *cis-* and *trans*-spliced introns for both species (7 and 14 intronic nucleotides respectively for *G. lamblia;* 11 and 21 intronic nucleotides respectively for the longer conserved consensus sequences of *S. salmonicida);* (ii) for each position within the boundary, each of the four nucleotides was assigned a score equal to the frequency of the nucleotide in known introns plus 0.05 (added to account for the possibility that newly found introns could use nucleotides not observed among the small sets of known introns); (iii) the raw score for each boundary for each potential case was calculated as the log of the product of the scores across sites; (iv) the final score was calculated as the maximum possible score minus the raw score (thus the maximum possible final score is zero). Scores were calculated for each position within five nucleotides downstream and upstream of the apparent junction, and the maximum among these scores was used as the score for the boundary. In addition, for *G. lamblia*, each potential cases of trans-splicing was analyzed by eye; for the larger number of potential cases for *S. salmonicida*, the cases with the top 500 scores were analyzed by eye. To determine evidence for trans-splicing in the various datasets, 12 RNA-seq datasets from Ansell et al. (2015) were downloaded from SRA (Accession PRJNA298647). The first 100 nucleotides of each read for the Franzen et al. and Ansell et al. datasets were mapped against the spliced and unspliced forms of each trans-spliced intron using Bowtie with default parameters, with reads that mapped to only the spliced form being taken as evidence for splicing.

